# Functional diversity across families of bacterial metalloregulators: what can we learn about specificity from sequence similarity?

**DOI:** 10.64898/2026.05.01.721428

**Authors:** Johnma J. Rondón, Giuliano T. Antelo, Daiana Capdevila

## Abstract

The exponential growth of sequence databases, driven by large-scale genome sequencing, has created a major challenge for the functional annotation of proteins, particularly within highly divergent families such as bacterial metal-responsive transcription factors. In these systems, low sequence identity places many proteins within the so-called “twilight zone” of the proteome, where homology inference and functional assignment become unreliable. Here, we integrate sequence similarity networks (SSNs) with structural approaches to explore the functional diversity of twelve metalloregulatory families. Using SSNs, we partition each family into putative isofunctional clusters and map available experimental annotations onto network topology, revealing substantial heterogeneity in both sequence diversity and functional characterization across families. While some families, such as ArsR and CsoR, display relatively well-defined functional landscapes, others, including LysR, TetR, and GntR, remain largely unexplored despite their large sequence space. Structural comparisons further show that, despite extensive sequence divergence, conserved architectural features underpin DNA recognition and regulatory mechanisms across families. Focusing on the MerR and Fur families, we identify conserved residues associated with inducer binding and DNA recognition, and uncover distinct functional subgroups, including metal-specific sensors and regulators with alternative signaling mechanisms. Finally, we demonstrate that cluster-derived HMM profiles enable sensitive detection of candidate regulators in non-model genomes, revealing lineage-specific expansions and diversification of metal-sensing repertoires. Together, our results provide a framework for mapping functional diversity in highly divergent protein families and highlight the potential of combining SSNs and structural information to guide the discovery of novel transcriptional sensors.

## INTRODUCTION

Metal ions play essential roles in bacterial physiology, acting as cofactors and signals for transcriptional responses under stress conditions. To maintain metal homeostasis, bacteria rely on specialized metalloregulatory transcription factors, including representatives of the ArsR, MerR, Fur, MarR, and other families (Capdevila et al. 2024; Chandrangsu et al. 2017). Although these regulators have been extensively studied in a handful of model organisms, the vast majority of their homologs across bacterial diversity remain uncharacterized. This gap limits functional annotation in comparative genomics and the ability to identify novel sensors in pathogens or environmental species.

Protein families are defined by their functional domains, which constitute the fundamental unit of evolution and can diverge at different rates across lineages (Forslund et al. 2019). During this process, families can expand through gene duplications, horizontal gene transfer (HGT), or de novo origin events, or contract through deletions and pseudogenization, even disappearing in certain lineages (Merhej et al. 2009). Domain duplication has been particularly frequent, reaching proportions close to 70% in prokaryotes and up to 90% in eukaryotes, which has favored their combination with different partners and given rise to the enormous diversity of multidomain proteins observed (Chothia et al. 2003). The exclusivity of a domain in a genome can increase its adaptive value and functional dependence, while domains whose functions can be replaced by others are more likely to be lost during evolution (Nasir et al. 2014; Koonin 2009). Thus, gain and loss events determine the dynamics of gene turnover within a family and shape protein repertoires across lineages.

The expansion of genomic sequencing has enabled the identification of thousands of new sequences belonging to diverse families, facilitating the detection of homologs across the tree of life and understanding how the functions evolve (Ellegren 2014). However, it has also revealed that a significant proportion of genes lack a known function, falling into the so-called “twilight zone,” characterized by low levels of sequence identity that hinder the inference of homology (Ponting 2017) or, in more extreme cases, we find sequences for which it is not possible to assign a function or a reliable evolutionary relationship by any known method; at that point we speak of the ‘dark matter’ of the proteome (Pavlopoulos et al. 2023). In this context, the detection of distant homologs requires more sensitive approaches than simple sequence comparison, where protein structure emerges as a more conserved evolutionary proxy, allowing the tracing of ancestral relationships even when the sequence signal has been almost completely lost (Orengo and Thornton 2005).

Indeed, despite the vast knowledge available in databases for many protein families, the reality is that we only know a small fraction of the repertoire present in nature. This is why Sequence Similarity Networks (SSNs) offer a scalable approach to visualizing and analyzing protein superfamilies with thousands of sequences, enabling the identification of putative functional clusters based on sequence divergence (Copp et al. 2018; Mai et al. 2016; Navarro-Muñoz et al. 2020). Integrating SSNs with curated annotations offers an opportunity to assess the current state of knowledge for each family, quantify the number of regulators with known inducers, and identify clusters that remain unexplored. Such quantitative assessments help prioritize families or clades for deeper functional investigation. Beyond descriptive analyses, SSNs can inform the construction of cluster-specific profile Hidden Markov Models (HMMs), which allow sensitive detection of candidate sensors in bacterial genomes (Sharma et al. 2016; Sinha et al. 2018). This is particularly relevant for non-model organisms, where gene annotation is often limited. By combining SSNs, HMM-based genomic mining, phylogenetics, and a structural approach, we aim to propose a unified framework for mapping the functional landscape of metalloregulatory families and for discovering previously unrecognized sensors, particularly in poorly explored genomes.

## Methods

### 1. Sequence similarity networks (SSNs) and diversity of metaloregulator families

SSNs were generated for twelve bacterial transcriptional regulator families involved in metal sensing (ArsR, MerR, MarR, Fur, DtxR, CsoR, CopY, NikR, Rrf2, LysR, GntR, and TetR) using the EFI– Enzyme Similarity Tool (Gerlt et al. 2015). Protein sequences were retrieved based on Pfam and InterPro family identifiers (Table S1), using database versions current as of January 2024. To reduce the inclusion of multidomain proteins and truncated sequences, family-specific sequence-length filters were applied before network construction. SSNs were generated across a range of alignment score (AS) thresholds, and the final AS cutoff for each family was selected based on the emergence of clusters consistent with putative isofunctional groups, based on available functional annotations. Networks were visualized and analyzed using Cytoscape v3.10.4 (Shannon et al. 2003).

### 2. Dataset selection, structural comparison across families, and estimation of intra-family sequence diversity

Representative three-dimensional structures were selected for each of the twelve transcription factor families from the PDB. For each family, a single structure representing the biologically relevant oligomeric assembly was chosen. Structures were manually curated to remove non-protein components, to focus comparisons on the protein scaffold and oligomeric architecture. Structural similarity was assessed using the DALI software (Holm 2022). DALI performs pairwise structural alignments by comparing intramolecular Cα–Cα distance matrices, enabling the detection of fold-level similarities independent of sequence identity. All-against-all structural comparisons were conducted among the oligomeric assemblies. Structural similarity was quantified using DALI Z-scores. The distance-based tree was customized using the iTOL web portal (Letunic and Bork 2021). To estimate average pairwise sequence identity within each family, random subsampling was performed using CD-HIT (Li and Godzik 2006). For each family, 5000 sequences were randomly selected from the filtered dataset. Exhaustive pairwise alignments within each subsample were computed using BLASTp (Camacho et al. 2009). Pairwise sequence identities were calculated for all sequence combinations within each subsample. This procedure was repeated across ten independent random replicates to account for sampling variance. The mean pairwise sequence identity across replicates was used as an estimate of overall sequence diversity for each family.

### 3. Functional evolution of the MerR and Fur families of metal-responsive transcriptional regulators

Based on the information derived from the SSN, clusters with putative functional annotation were analyzed. For each cluster, protein sequences were aligned using MAFFT v7.505 (Katoh and Standley 2013) and cleaned with TrimAl v1.5 (Capella-Gutiérrez et al. 2009). Sequence logos were generated with WebLogo (Crooks et al. 2004) to identify conserved motifs potentially associated with functional specificity in each cluster. In addition, sequences of previously characterized MerR and Fur regulators were included to explore their phylogenetic relationships across SSN-defined clusters. To this end, we used a structure-based approach with the FoldTree server (Moi et al. 2025) employing structural models predicted with AlphaFold3 (Abramson et al. 2024).

### 4. Identification of metalloregulator repertoires in non-model bacteria using SSN-derived information

We used the information derived from isofunctional clusters associated with metal sensing in the ArsR, MerR, and Fur families to guide a targeted search in non-model bacterial genomes. We selected well-characterized reference organisms such as *Mycobacterium tuberculosis* H37Rv and *Vibrio cholerae* RFB16, along with their less-characterized counterparts, *M. avium hominissuis* (GCA_004345205.2) and *V. vulnificus* NBRC 15645. Hidden Markov Model (HMM) profiles were constructed for each isofunctional cluster using HMMER (Moi et al. 2025; Eddy 2011) and were then used to identify candidate genes in bacterial genome assemblies through the BITACORA pipeline (Vizueta et al. 2020). Searches were performed for clusters associated with different metal inducers (zinc, mercury, nickel, and lead) and under two stringency thresholds (E-values of 1e−3 and 1e−20), representing relaxed and stringent homology criteria, respectively. The resulting gene repertoires were compared across related species, and their chromosomal distributions were analyzed relative to the origin of replication, approximated by the genomic position of the *dnaA* gene. Finally, candidate metal sensors were compared with structurally characterized regulators to assess their potential functional assignments.

## RESULTS AND DISCUSSION

### 1. Structural diversity of twelve families of bacterial metalloregulators

Transcriptional regulator families in bacteria have undergone remarkable diversification at multiple levels, ranging from their sequence and architectures to their regulatory mechanisms. These regulators can function as transcriptional repressors, activators, or, in certain contexts, both (Fig. S1A), while still conserving key structural elements required for DNA recognition and regulatory activity, such as variants of the helix–turn–helix (HTH) domain necessary for DNA binding. The deep diversification of many regulator families across bacterial lineages has largely erased detectable sequence homology among members of the same family; proteins in this scenario are considered to be in the twilight zone. Phylogenetic methods based solely on sequence information often lack the robustness required to infer evolutionary relationships among these families and are even more limited in tracing their ancestral origins. For this reason, and taking advantage of advances in protein structures prediction, it is increasingly common to incorporate structural information to investigate evolutionary relationships between and within protein families (Abramson et al. 2024; Moi et al. 2025). Although a common evolutionary ancestor among bacterial metalloregulator families is no longer detectable at the sequence level, we employed a global structural distance approach to compare representative members from the twelve families analyzed (Fig. 1).

**Fig. 1.**
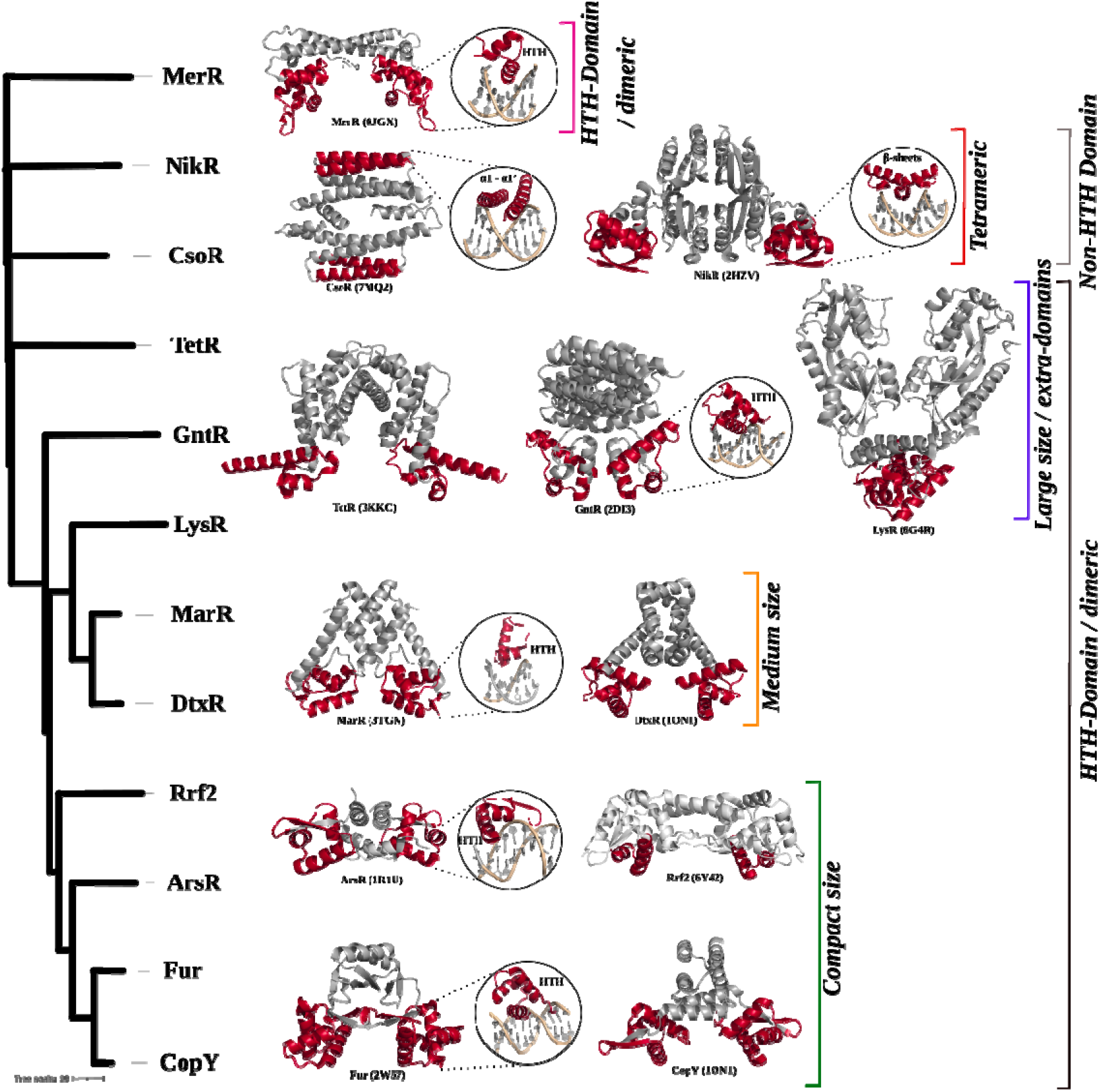
Relationships among metalloregulator families. Structural similarity dendrogram generated from pairwise structural comparisons (Z-scores) calculated using the DALI server. The analysis was performed on representative oligomeric assemblies (PDB structures) from each family. Selected structures are shown alongside the dendrogram, with the DNA-binding domain highlighted in red. PDBs used: MerR JGX; NikR 2HZV; CsoR 7MQ2 TetR 3KKC; GntR 2DI3; LysR 6G4R; MarR 3TGN; DtxR 1ON1; Rrf2 6Y42; ArsR 1R1U; Fur 2W57; CopY 1OKR.

Structural comparison based on internal distances derived from three-dimensional geometry between representatives of each family revealed three main structural groups: i) regulators with an HTH domain, ii) non-HTH regulators, and iii) the MerR family. Within the first group, we observed a wide diversity of regulators sharing the DNA-binding HTH domain, which can be subdivided into three subgroups according to their size and structural complexity (Fig. S1B). The first subgroup corresponds to compact conformations, such as those present in the ArsR and Fur families, as well as Fur and CopY, which also incorporate an extended dimerization domain via a linker. The second subgroup includes medium-sized regulators, such as the MarR and DtxR families, in which the dimerization domain exhibits greater structural complexity. Finally, the third subgroup comprises the larger HTH regulators, belonging to the GntR, TetR, and LysR families, whose dimerization domains are even more extensive and, in many cases, in addition to participating in inducer recognition, may also have other not necessarily regulatory functions. The second major group corresponds to the non-HTH regulators, which, besides interacting with DNA through alternative structural motifs, are characterized by their tetrameric oligomeric organization, either homotetrameric (as in CsoR) or heterotetrameric (as in NikR), in contrast to the other families, which are predominantly homodimeric. In particular, CsoR binds to DNA at a single site via the α1 and α1’ helices, while NikR uses β-sheets that insert into the major groove of the DNA. The third group is represented by the MerR family, which, although it possesses an HTH domain, differs from the others because it has a relatively simple dimerization domain. In contrast, the DNA-binding module exhibits greater structural complexity, probably associated with its characteristic function as a transcriptional activators that induce DNA distortion (Fig. 1). A common feature across families is the organization of their functional components. In general, they exhibit an arrangement in which the N-terminal end is specialized for DNA binding, while the C-terminal end contains domains responsible for dimerization and, in many cases, for inducer binding. Furthermore, dimer formation appears to be the norm, implying in many cases the need for two DNA-binding sites separated by a spacer, whose length varies among the different families.

Differences in structure and sequence between families are evident; their functional diversity in terms of the chemical species that they respond to, and their degree of functional overlap are still unclear. To evaluate the functional diversity within metalloregulator families, we constructed sequence similarity networks (SSNs) for each family, using data available in UniProt and InterPro databases (Table S1). Then, depending on the availability of experimentally characterized regulators, we mapped known regulators onto specific network clusters. Using this approach, we identified putative clusters associated with specific metal responses (Zn, Ni, Cu, Hg, and As ions), clusters exhibiting apparent inducer promiscuity, and clusters specialized in sensing reactive sulfur sulfane species (RSS) or reactive oxygen species (ROS) across distinct regulatory families.

The SSN clusters containing one or more proteins responsive to the same inducer were classified as putative isofunctional groups, under the assumption that sequence similarity at the selected threshold reflects shared functional properties. To integrate family size, functional annotation, and network topology, we summarized for each family: i) total of sequences, ii) total of clusters, iii) clusters containing experimentally characterized regulators, and iv) sequences in clusters containing experimentally characterized regulators (Fig. 2A). This comparative overview highlights striking differences among families, for instance, some of them are both large and functionally diversified, whereas others are comparatively compact and dominated by a single characterized regulatory mode. Importantly, family size alone does not predict the degree of functional annotation: large families such as LysR, TetR, GntR, and MarR contain many sequences, but they have relatively few clusters with experimentally characterized metal-responsive members. This observation could be explained either because metal regulation is only a small part of the regulatory landscape for these families, or it is simply a lack of sampling or a gap in their characterization. In contrast, families such as ArsR, CsoR, and NikR display a higher proportion of clusters with known functional assignments relative to their total sequence diversity, suggesting greater functional completeness within the currently annotated space.

**Figure 2.**
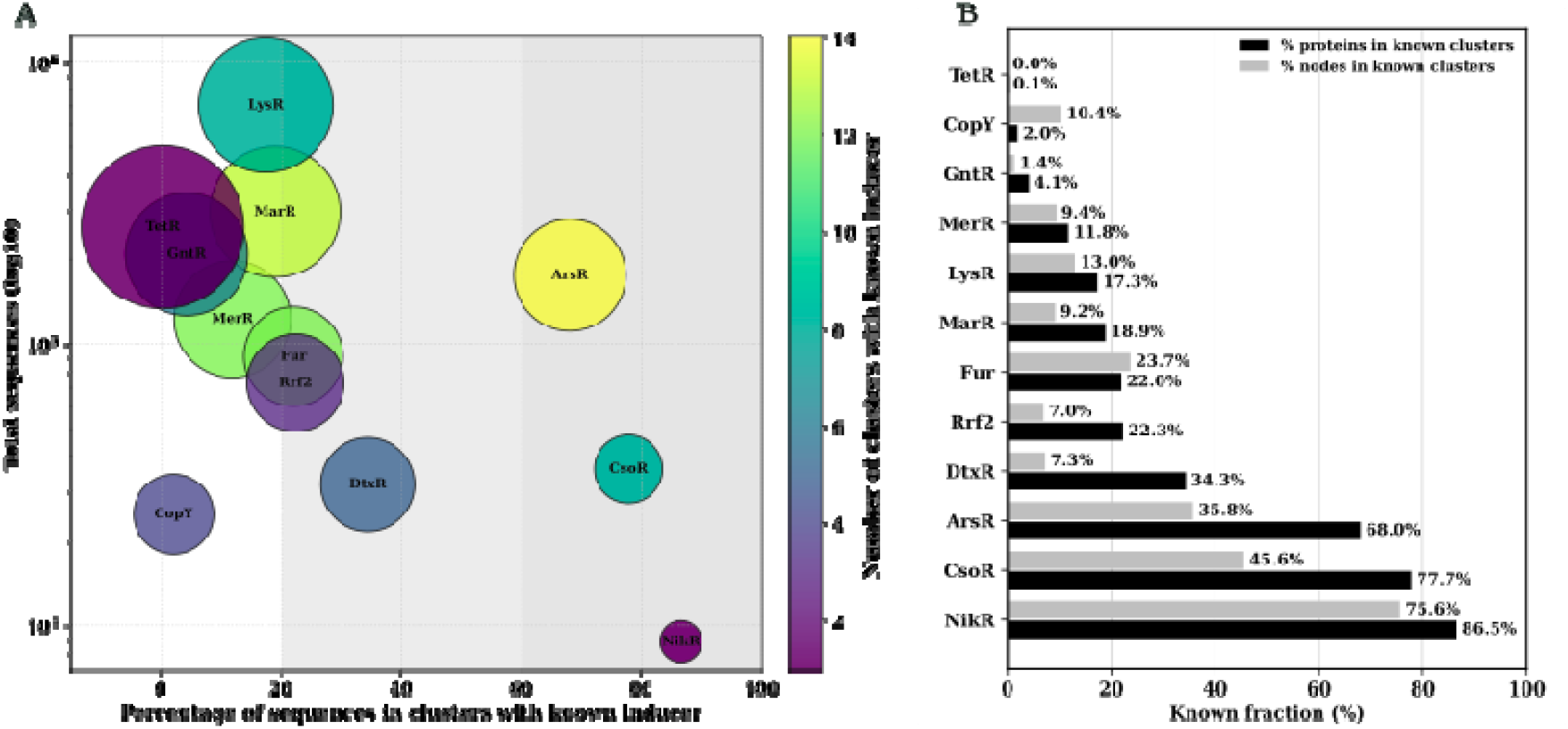
Sequence clustering of 12 metalloregulatory protein families using SSNs generated with the EFI-EST tool. **A**. Bubble plot showing the relationship between family size (total number of sequences; y-axis) and family diversity (total number of clusters; bubble size). Families are classified into *poorly, partially*, and *well* characterized groups based on the proportion of clusters containing proteins with known inducers (x-axis); the background color bands illustrate these categories. The color of each bubble represents the number of clusters with experimentally characterized members, providing an additional measure of the functional diversity known for each family. **B**. Discriminatory power of sequence similarity networks (SSNs) across 12 regulatory families. Bars indicate the fraction of proteins (black) and nodes (gray) that map to SSN clusters containing at least one experimentally characterized member. Such clusters are treated as functionally known, providing putative functional information for the remaining sequences within each cluster. For the TetR, GntR, LysR, and MarR families, values correspond only to the metal-responsive subgroups.

Across families, this approach identified between one and fourteen clusters, highlighting substantial variability in functional diversification, although in some cases this variation may reflect the intrinsic specificity of the family (Fig. 2A). For instance, in the NikR family, we identified a single dominant cluster encompassing the majority of sequences (86.5% of the total). Given that this family is well established as a highly specialized nickel sensor, it is reasonable to infer that this cluster predominantly responds to nickel as its inducer. Therefore, the NikR family shows a more complete functional landscape than other families, in which the fraction of sequences in clusters with known function is much lower, assuming that all sequences within a cluster share a common function. In families such as TetR, CopY, and GntR, the percentage of sequences in known-function clusters is below 5% (Fig. 2B). While this clustering pattern is influenced by the alignment score threshold used during SSN construction, the level of network fragmentation could be refined as more proteins with known inducers are identified within each family. Overall, this SSN-based approach provides a framework for partitioning each family into putative functional subfamilies, beyond serving as a strategy for functional classification, this framework can also guide efforts to discover novel sensors responsive to specific inducers of interest, thereby offering a roadmap for identifying new proteins with potential biotechnological applications, for example in biosensor development.

Although the literature reports that families of metalloregulatory proteins are highly diverse, there are currently no quantitative assessments of this diversity. We estimated average pairwise sequence identity within each family by random subsampling (5,000 sequences per family) followed by exhaustive BLASTp comparisons. Across the twelve families, average sequence identity ranged from 31% to 50%, confirming substantial divergence within most groups (Fig. S2; Table S1). Families such as ArsR, CopY, Fur, MerR, and Rrf2 exhibited particularly low average identities (<38%), consistent with extensive sequence diversification and the presence of multiple SSN clusters. In contrast, CsoR, DtxR, MarR, and NikR showed comparatively higher conservation (>43%), in agreement with more compact network topologies and fewer functionally distinct clusters. It is important to note that identity estimates for GntR, LysR, and TetR were derived from selected subgroups rather than entire superfamilies and therefore underestimate their full sequence diversity. Families that combine high sequence diversity with multiple isofunctional clusters represent particularly promising systems for investigating the emergence of new metal-sensing specificities.

### 2. The MerR family can be separated into two groups: canonical metalloregulators and non-metalloregulators

The HTH MerR family of transcriptional regulators exhibits remarkable diversity in both the types of inducers recognized and regulatory mechanisms. The classical mechanism described for this family, particularly among metal-sensing regulators, involves binding to the operator and inducing an inducer-dependent conformational change that promotes DNA underwinding and torsion, thereby realigning the -35 and -10 elements of suboptimal σ^70^ promoters and facilitating transcriptional initiation. Binding sites in these promoters typically consist of an inverted repeat separated by a spacer of approximately 19–20 bp, which is longer than the optimal spacing required for transcription initiation. In the absence of an inducer, MerR-family proteins bind as dimers to this operator region through their N-terminal HTH domain, positioning the α2 recognition helices within adjacent major grooves of the DNA. In this context, the logos analysis showed a conserved IRYYExxG motif, which may contribute to base-specific recognition of the operator (Fig. 3C). Upon inducer binding, conformational changes transmitted through the coiled-coil dimerization helix generate torsional strain in the bound DNA, effectively underwinding the promoter and realigning the -35 and -10 elements into a geometry compatible with σ^70^ recognition. However, this mechanism is not universal within the family. Indeed, in addition to acting as transcriptional activators through structural remodeling of DNA, some members can function as repressors, either by sterically blocking access of RNA polymerase or through alternative regulatory mechanisms, such as a co-repressor.

**Figure 3.**
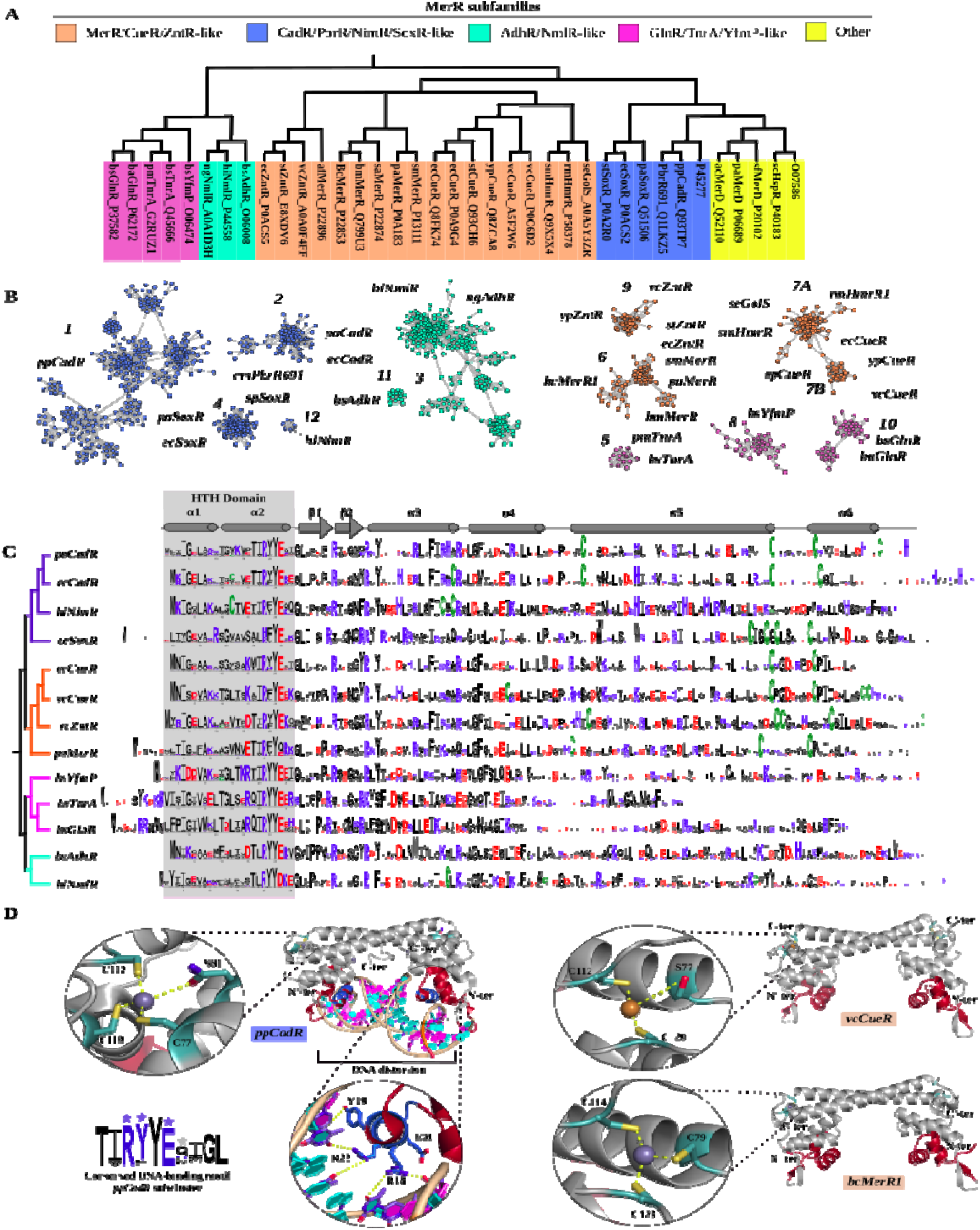
Functional and structural diversification of the MerR family. A) Structural clustering of curated sequences from different MerR subfamilies was performed using FoldTree. B) Functional clusters identified in the MerR family SSN that contain experimentally characterized members. Structurally associated groups primarily linked to metal sensing are highlighted in violet and orange clusters, as well as groups associated with responses to organic compounds and redox stress (cyan and magenta clusters). C) Sequence logos corresponding to each cluster, highlighting the N-terminal region containing the DNA-binding domain (colored background), allowing comparison of the conservation of the DNA-recognition module across functional subfamilies. D) Representative three-dimensional structures of metal-sensing regulators (ppCadR: PDB 6JNI; vcCueR: AF3 model + Cu^2^□; bcMerR: AF3 model + Zn^2^□) and non-metal sensors (hiNmlR: PDB 5D8C), illustrating structural differences associated with inducer type.

This functional diversity reflects broad structural plasticity within the MerR family. We focused primarily on the group referred to as “canonical” MerR regulators, which includes proteins with the classical structural architecture (an N-terminal HTH domain, a coiled-coil dimerization helix, and a compact C-terminal sensor domain) and which, in most cases, respond to metal ions through the DNA-torsion activation mechanism. Indeed, several canonical MerR sensors can respond to drugs and toxic compounds, but those were excluded from this analysis (length filtering). Consequently, our study centers on a structurally coherent subset of compact MerR regulators.

The high degree of divergence among MerR subfamilies, a situation comparable to that observed in other large transcriptional regulator families, the reconstruction of evolutionary relationships based exclusively on sequence information, even when restricted to the DNA-binding HTH domain, is poorly robust and yields low statistical support. For this reason, we complemented our analysis with structure-based approaches. Although structural similarity does not allow the establishment of direct evolutionary relationships with the same rigor as a classical phylogenetic analysis, it provides valuable information regarding functional convergence and architectural conservation. For this purpose, we used FoldTree, which relies on the Foldseek algorithm and a structural alphabet (3Di) that transforms three-dimensional information into a representation of local structural states, in contrast to DALI, which compares proteins by evaluating global architectural similarity through structural alignment. Whereas DALI is particularly robust for detecting global fold similarities (including between families that are not necessarily homologous), FoldTree is useful for exploring finer structural relationships within curated sets of related proteins (subfamilies). Applying FoldTree to structural representatives curated from InterPro for each MerR subfamily, we identified two main groups (Fig. 3A): one composed predominantly of metal-ion-responsive sensors and another grouping non-metal regulators. The clade including CadR (Cd^2+^), NimR (Ni^2+^), CueR (Cu^2+^), ZntR (Zn^2+^), and MerR (Hg^2+^) consists exclusively of classical metal-sensing regulators. Notably, this group also contains SoxR, a redox sensor dependent on [2Fe–2S] centers, suggesting that, despite responding to different inducers, these regulators share a conserved structural core associated with the DNA-torsion activation mechanism. In contrast, the clades corresponding to NmlR and AdhR, although often considered canonical, show closer structural proximity to the non-canonical regulators GlnR and TnrA, which do not use metal ions as inducers and are instead involved in nitrogen metabolism responses. This indicates that structural architecture may reflect bigger mechanistic differences than traditional functional classifications suggest.

The structural clustering obtained is consistent with the clustering derived from sequence similarity in the SSNs constructed for the family (Fig. 3B). This concordance suggests that, when appropriate construction and filtering criteria are applied, SSNs can recover functional subfamilies with a considerable degree of structural accuracy. A particularly evident difference among metal-sensing MerR subfamilies is the conservation of reactive cysteines that are key for metal ion coordination. In contrast, these cysteines are absent in subfamilies such as TnrA, GlnR, AdhR, and NmlR (Fig. 3C). In classical metal-sensing regulators, metal coordination typically involves at least two cysteines incorporated into the motif CX_7-8_C C-terminal, arranged in a geometry compatible with trigonal or tetrahedral binding, triggering the allosteric change responsible for DNA torsion and transcriptional activation (Fig. 3D).

### 3. The Fur family is strongly oriented towards metal regulation, albeit presenting higher metal promiscuity

We next examined in detail the Ferric uptake regulator (Fur) family, a superfamily of bacterial metalloregulators that has been reported to sense Mn^2^□, Co^2^□, Ni^2^□, Zn^2^□, Fe^2^□, as well as H_2_O_2_ (Capdevila et al. 2024). Although the best-characterized members of this family use Fe^2^□ as the cognate metal in their regulatory allosteric site, different subfamilies have evolved alternative metal specificities, potentially leading to sensing crosstalk among related regulators. Structure-based comparisons reveal four main groups within the Fur superfamily (Fig. 4A), corresponding to regulators associated with Fe^2^□ and Mn^2^□ sensing (Fur/Mur-like), Zn^2^□ and Ni^2^□ sensing (Zur/Nur-like), oxidative stress sensors such as PerR, and heme-dependent regulators such as Irr. The first group includes the classical Fur/Mur sensors, predominantly associated with iron and manganese metabolism. The close structural relationship between Fur and Mur suggests that specificity toward Fe^2^□ or Mn^2^□ may have emerged through subtle modifications of a common ancestral scaffold, reflecting the physiological interconnection between iron and manganese homeostasis. The second group corresponds to the Zur/Nur subfamily. In Zur regulators, Zn^2^□ plays a dual role: it may act as a structural component stabilizing the dimer and also serve as the cognate regulatory metal. Sequence logo analysis indicates reduced conservation of histidines within the HHxH motif relative to Fur/Mur proteins, whereas the cysteines of the Cx□C motif remain highly conserved, consistent with a structural metal-binding site based on sulfur coordination. Within this group, Nur represents a specialization toward Ni^2^□ sensing while maintaining an overall similar structural architecture. The third group comprises PerR-like regulators, which function as oxidative stress sensors. In these proteins, the metal ion (Fe^2^□ or Mn^2^□) does not simply serve a structural or allosteric role but directly participates in redox reactions that lead to oxidation of coordinating residues, triggering conformational changes that modulate DNA-binding affinity. Finally, the fourth group corresponds to Irr, a heme-dependent regulator involved in the control of iron metabolism. Unlike the other subfamilies, Irr uses heme as the signaling molecule, thereby integrating iron availability with cellular metabolic status. The close evolutionary relationship among Fur regulators sensing different metal ions has resulted in the emergence of multiple metal coordination sites with distinct functional roles. Some of these sites serve structural purposes, whereas others participate directly in allosteric regulation. Members of the Fur family typically contain between two and three metal-binding sites per monomer (i.e., four to six sites per dimer), although not all of them function as regulatory sites. Among these, the so-called site 2 corresponds to the functional allosteric site and is formed by conserved combinations of histidines, cysteines, and acidic residues, frequently organized in motifs of the type HHxHx□Cx□C. This motif is broadly conserved across subfamilies (Fig. 4C) and can adopt different coordination geometries, with tetrahedral coordination being the most common (Fig. 4D). Sequence similarity network (SSN) clustering reveals a number of functionally characterized clusters comparable to that observed for the MerR family, although with more extreme cluster sizes. In particular, each subfamily contains at least one very large cluster accompanied by several much smaller clusters. While functional separation is generally well resolved, the high sequence similarity between Fur and Mur regulators remains one of the most difficult relationships to resolve within the network (Fig. 4B). With respect to DNA recognition, Fur-family regulators contain an N-terminal winged helix–turn–helix (wHTH) DNA-binding module. In this region, helices α3 and α4 are in proximity to the DNA major groove, and the conserved TVYRTLQ motif likely contributes to the specificity for Fur-box operator sequences (Fig. 4D).

**Figure 4.**
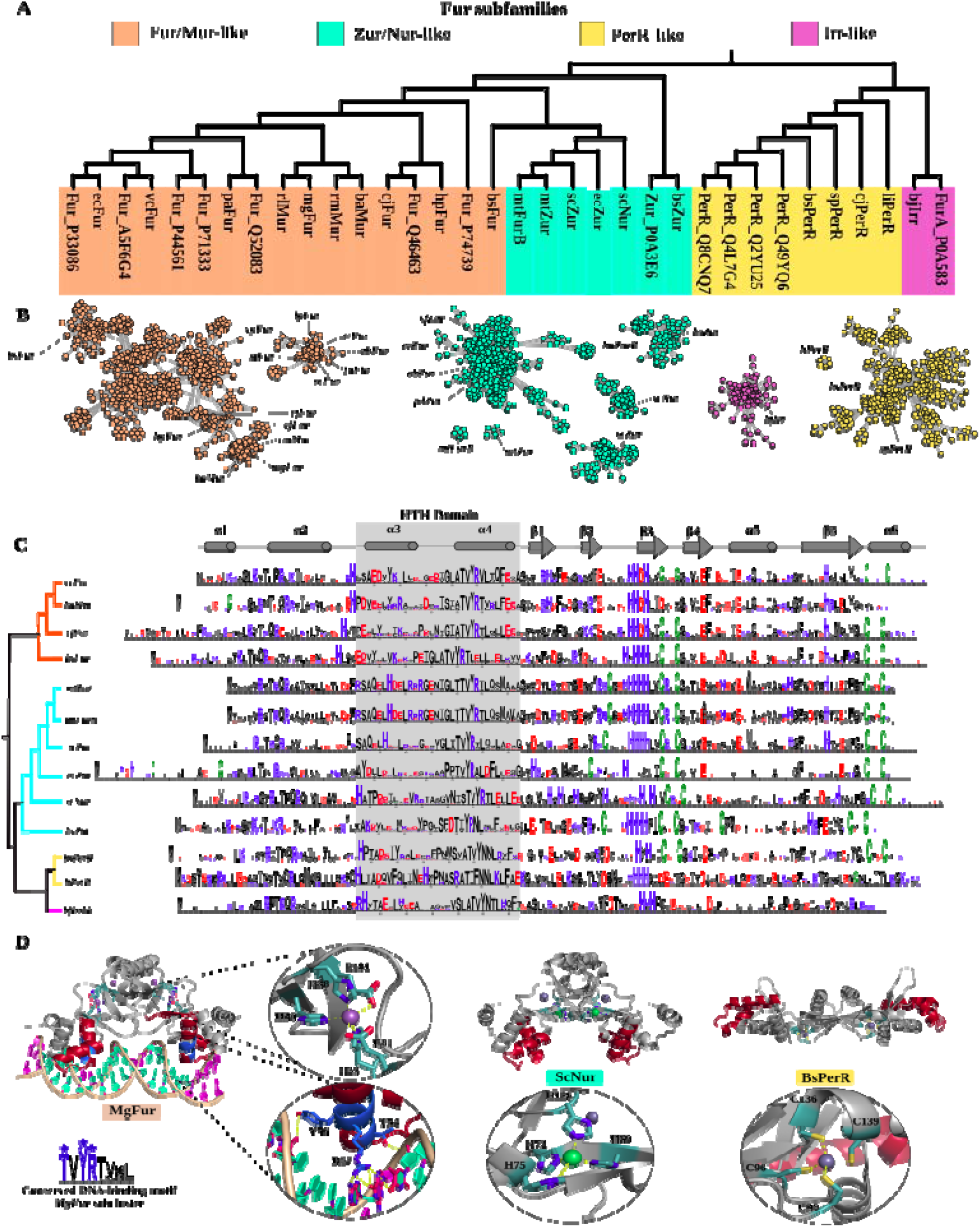
Functional and structural diversification of the Fur family. A) Structural clustering of curated sequences from different Fur subfamilies was performed using FoldTree. B) Functional clusters identified in the Fur family SSN that contain experimentally characterized members. Structurally associated groups primarily linked to metal sensing are highlighted in red, cyan, and magenta colors, and clusters associated with redox responses are shown in yellow. C) Sequence logos corresponding to each cluster, highlighting the N-terminal region containing the DNA-binding domain (colored background), allowing comparison of the conservation of the DNA-recognition module across functional subfamilies. D) Representative three-dimensional structures of metal-sensing regulators (ppCadR: PDB 6JNI; vcCueR: AF3 model + Cu^2^□; bcMerR: AF3 model + Zn^2^□) and non-metal sensors (hiNmlR: PDB 5D8C), illustrating structural differences associated with inducer type.

### 4. Functional clustering derived from SSNs facilitates the identification of metalloregulator repertoires and novel sensors in non-model genomes

HMM profile–guided homology searches provide a highly sensitive strategy for detecting distant members of protein families, even when sequence identity falls below 30%. In this study, we leveraged previously defined functional clustering of metalloregulator families to guide the identification of putatively metal-specific sensors across bacterial genomes with different levels of characterization. To this end, we selected subfamilies associated with Zn^2^□ (CzrA-ArsR, ecZntR-MerR, bsZur-Fur, and ecZur-Fur), Hg^2^□ (pmMerR-ArsR and paMerR-MerR), Ni^2^□ (mtNmtR-ArsR, hiNmlR-MerR, and scNur-Fur), and Pb^2^□ (PbrR-MerR) sensing. Both HMM profiles and manually curated representative sequences were used to guide searches in two reference genomes, *M. tuberculosis* H37Rv and *V. cholerae* RFB16, as well as in two phylogenetically related but less characterized species, *M. avium hominissus* and *V. vulnificus* NBRC15645. Overall, the BITACORA-based pipeline enabled the recovery of candidate protein repertoires for each subfamily, allowing comparisons both at the regulatory family level and in terms of metal specificity. Under a relaxed threshold (E-value = 1e−3), the number of retrieved proteins provided an approximation of the total repertoire of family members present in each genome, including more divergent homologs or candidates whose functional specificity cannot yet be confidently resolved. Notably, this approach is inherently biased toward metalloregulators and may overlook members of these families that respond to non-metallic inducers. In this context, the highest abundances were generally observed for ArsR and MerR subfamilies, whereas members of the Fur family tended to occur in more limited numbers. For instance, in *M. tuberculosis*, ArsR-type subfamilies associated with Zn^2^□ and Ni^2^□ displayed expanded repertoires, with approximately 11–14 candidates, whereas in *V. cholerae*, the corresponding range was lower, approximately 3–9 proteins (**Fig 5A**). These observations suggest that metal-responsive regulatory repertoires may be more expanded in mycobacterial lineages compared to vibrios. Applying a more stringent threshold (E-value = 1e−20) resulted in a marked reduction in the number of retained proteins across nearly all subfamilies. This behavior is consistent with the expectation that stricter criteria preferentially retain homologs with higher similarity to the reference isofunctional clusters, and therefore a higher likelihood of corresponding to functionally specialized metal sensors. Thus, while the relaxed threshold provides an estimate of overall family diversity, the stringent threshold could define a subset of higher-confidence candidates. This trend is consistently observed across all metal categories and supports the combined use of both thresholds to distinguish between global family diversity and putatively metal-specific regulators.

**Figure 5.**
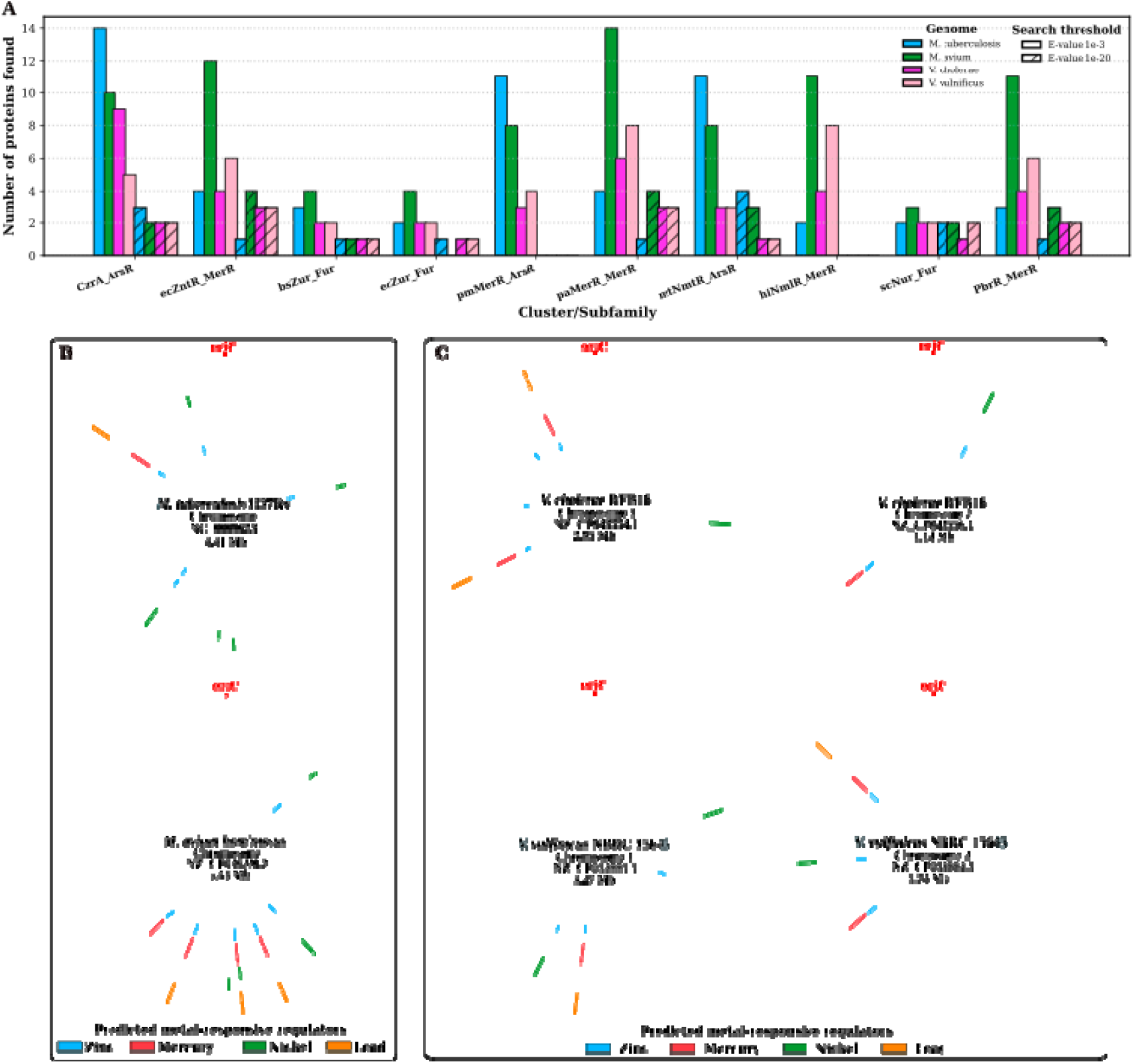
Identification of candidate metalloregulators from the ArsR, MerR, and Fur families across bacterial genomes. (A) Number of candidate metalloregulatory genes identified using HMM profiles derived from isofunctional clusters associated with Zn^2^□, Hg^2^□, Ni^2^□, and Pb^2^□ sensing. Searches were compared across genomes with different levels of annotation, including *M. tuberculosis* and *M. avium*, as well as *V. cholerae* and *V. vulnificus*, using two significance thresholds (E-values of 1e−3 and 1e−20). (B–C) Relative chromosomal distribution of candidate metalloregulator genes in mycobacterial and vibrios genomes, respectively. Track colors indicate the metal specificity associated with each identified subfamily, while the length of each strip reflects the number of subfamilies detected for a given metal; strips spanning the entire track correspond to a single identified subfamily. (D) AlphaFold3 structural models of selected candidate proteins identified in the less-characterized genomes *M. avium* and *V. vulnificus*.

Metal-specific analyses further revealed lineage-dependent differences. For Zn^2^□ associated sensors, ArsR and MerR subfamilies exhibited notable expansion, particularly in *M. tuberculosis* and *M. avium*, whereas Fur subfamilies showed more conserved copy numbers across species, typically ranging from one to a few candidates under stringent conditions. In the case of Hg^2^□, a marked disparity was observed between genera, with ArsR-type subfamilies being more abundant in mycobacteria; however, under stringent criteria, only the paMerR-MerR subfamily retained robust candidates, suggesting stronger functional conservation in this group. Similarly, Ni^2^□ associated candidates from mtNmtR-ArsR and hiNmlR-MerR were more prevalent in mycobacterial genomes, whereas Fur-type Ni sensors were less represented (**Fig 5A**). Finally, the Pb^2^□ associated PbrR-MerR subfamily showed interspecies variation, with a more prominent presence in the less-characterized genomes (*M. avium* and *V. vulnificus*) compared to their better-studied counterparts. Altogether, these observations indicate that the composition of metalloregulator repertoires is shaped not only by structural family constraints but also by lineage-specific evolutionary trajectories and ecological diversification of metal-sensing systems.

## CONCLUSIONS

In summary, this work demonstrates that the functional landscape of bacterial metalloregulators is shaped by a complex interplay between sequence divergence, structural conservation, and lineage-specific evolutionary trajectories. By integrating SSNs with structural analyses, we show that it is possible to resolve putative functional subfamilies even within the twilight zone of sequence similarity, linking sequence space to regulatory specificity. Our results highlight that family size does not necessarily correlate with functional characterization, revealing large unexplored regions of the proteome that likely harbor novel sensing mechanisms. Furthermore, the application of cluster-specific HMM profiles provides a powerful strategy to uncover candidate regulators in poorly annotated genomes, enabling comparative analyses of metal-sensing repertoires across taxa. Altogether, this framework not only advances our understanding of the evolution and diversification of metalloregulatory systems, but also establishes a scalable approach for the identification of new transcriptional sensors with potential applications in microbiology, biotechnology, and environmental monitoring.

## REFERENCES

Abramson, Josh, Jonas Adler, Jack Dunger, et al. 2024. “Accurate Structure Prediction of Biomolecular Interactions with AlphaFold 3.” Nature 630 (8016): 493–500.

Camacho, Christiam, George Coulouris, Vahram Avagyan, et al. 2009. “BLAST+: Architecture and Applications.” BMC Bioinformatics 10 (December): 421.

Capdevila, Daiana A., Johnma J. Rondón, Katherine A. Edmonds, Joseph S. Rocchio, Matias Villarruel Dujovne, and David P. Giedroc. 2024. “Bacterial Metallostasis: Metal Sensing, Metalloproteome Remodeling, and Metal Trafficking.” Chemical Reviews, ahead of print, December 10. 10.1021/acs.chemrev.4c00264.

Capella-Gutiérrez, Salvador, José M. Silla-Martínez, and Toni Gabaldón. 2009. “trimAl: A Tool for Automated Alignment Trimming in Large-Scale Phylogenetic Analyses.” Bioinformatics 25 (15): 1972–1973.

Chandrangsu, Pete, Christopher Rensing, and John D. Helmann. 2017. “Metal Homeostasis and Resistance in Bacteria.” Nature Reviews Microbiology 15 (6): 338–350.

Chothia, Cyrus, Julian Gough, Christine Vogel, and Sarah A. Teichmann. 2003. “Evolution of the Protein Repertoire.” Science (New York, N.Y.) 300 (5626): 1701–1703.

Copp, Janine N., Eyal Akiva, Patricia C. Babbitt, and Nobuhiko Tokuriki. 2018. “Revealing Unexplored Sequence-Function Space Using Sequence Similarity Networks.” Biochemistry 57 (31): 4651–4662.

Crooks, Gavin E., Gary Hon, John-Marc Chandonia, and Steven E. Brenner. 2004. “WebLogo: A Sequence Logo Generator.” Genome Research 14 (6): 1188–1190.

Eddy, Sean R. 2011. “Accelerated Profile HMM Searches.” PLoS Computational Biology 7 (10): e1002195.

Ellegren, Hans. 2014. “Genome Sequencing and Population Genomics in Non-Model Organisms.” Trends in Ecology & Evolution 29 (1): 51–63.

Forslund, Sofia K., Mateusz Kaduk, and Erik L. L. Sonnhammer. 2019. “Evolution of Protein Domain Architectures.” Methods in Molecular Biology (Clifton, N.J.) 1910: 469–504.

Gerlt, John A., Jason T. Bouvier, Daniel B. Davidson, et al. 2015. “Enzyme Function Initiative-Enzyme Similarity Tool (EFI-EST): A Web Tool for Generating Protein Sequence Similarity Networks.” Biochimica et Biophysica Acta 1854 (8): 1019–1037.

Holm, Liisa. 2022. “Dali Server: Structural Unification of Protein Families.” Nucleic Acids Research 50 (W1): W210–W215.

Katoh, Kazutaka, and Daron M. Standley. 2013. “MAFFT Multiple Sequence Alignment Software Version 7: Improvements in Performance and Usability.” Molecular Biology and Evolution 30 (4): 772–780.

Koonin, Eugene V. 2009. “Evolution of Genome Architecture.” The International Journal of Biochemistry & Cell Biology 41 (2): 298–306.

Letunic, Ivica, and Peer Bork. 2021. “Interactive Tree Of Life (iTOL) v5: An Online Tool for Phylogenetic Tree Display and Annotation.” Nucleic Acids Research 49 (W1): W293–W296.

Li, Weizhong, and Adam Godzik. 2006. “Cd-Hit: A Fast Program for Clustering and Comparing Large Sets of Protein or Nucleotide Sequences.” Bioinformatics (Oxford, England) 22 (13): 1658–1659.

Mai, Te-Lun, Geng-Ming Hu, and Chi-Ming Chen. 2016. “Visualizing and Clustering Protein Similarity Networks: Sequences, Structures, and Functions.” Journal of Proteome Research 15 (7): 2123–2131.

Merhej, Vicky, Manuela Royer-Carenzi, Pierre Pontarotti, and Didier Raoult. 2009. “Massive Comparative Genomic Analysis Reveals Convergent Evolution of Specialized Bacteria.” Biology Direct 4 (April): 13.

Moi, David, Charles Bernard, Martin Steinegger, Yannis Nevers, Mauricio Langleib, and Christophe Dessimoz. 2025. “Structural Phylogenetics Unravels the Evolutionary Diversification of Communication Systems in Gram-PositiveBacteria and Their Viruses.” Nature Structural & Molecular Biology 32 (12): 2492–2502.

Nasir, Arshan, Kyung Mo Kim, and Gustavo Caetano-Anollés. 2014. “Global Patterns of Protein Domain Gain and Loss in Superkingdoms.” PLOS Computational Biology 10 (1): e1003452.

Navarro-Muñoz Jorge C., Nelly Selem-Mojica, Michael W. Mullowney, et al. 2020. “A Computational Framework to Explore Large-Scale Biosynthetic Diversity.” Nature Chemical Biology 16 (1): 60–68.

Orengo, Christine A., and Janet M. Thornton. 2005. “Protein Families and Their Evolution-a Structural Perspective.” Annual Review of Biochemistry 74: 867–900.

Pavlopoulos, Georgios A., Fotis A. Baltoumas, Sirui Liu, et al. 2023. “Unraveling the Functional Dark Matter through Global Metagenomics.” Nature 622 (7983): 594–602.

Ponting, Chris P. 2017. “Biological Function in the Twilight Zone of Sequence Conservation.” BMC Biology 15 (1): 71.

Shannon, Paul, Andrew Markiel, Owen Ozier, et al. 2003. “Cytoscape: A Software Environment for Integrated Models of Biomolecular Interaction Networks.” Genome Research 13 (11): 2498–2504.

Sharma, Ronesh, Shiu Kumar, Tatsuhiko Tsunoda, Ashwini Patil, and Alok Sharma. 2016. “Predicting MoRFs in Protein Sequences Using HMM Profiles.” BMC Bioinformatics 17 (19): 504.

Sinha, Swati, Birgit Eisenhaber, and Andrew M. Lynn. 2018. “Predicting Protein Function Using Homology-Based Methods.” Bioinformatics: Sequences, Structures, Phylogeny, 289–305.

Vizueta, Joel, Alejandro Sánchez-Gracia, and Julio Rozas. 2020. “Bitacora: A Comprehensive Tool for the Identification and Annotation of Gene Families in Genome Assemblies.” Molecular Ecology Resources 20 (5): 1445–1452.

