## Supplementary material for "Functional diversity across families of bacterial metalloregulators: what can we learn about specificity from sequence similarity?"

Supplementary Information

Johnma J. Rondón^1,2^, Giuliano T. Antelo^1^, Daiana Capdevila^1^.

^1^ Fundación Instituto Leloir, Instituto de Investigaciones Bioquímicas de Buenos Aires (IIBBA-CONICET), C1405 Ciudad Autónoma de Buenos Aires, Buenos Aires, Argentina.

^2^ Departamento de Fisiología, Biología Molecular y Celular, Facultad de Ciencias Exactas y Naturales, Universidad de Buenos Aires, C1428 Ciudad Autónoma de Buenos Aires, Buenos Aires, Argentina.

Index figures and tables

- Figure S1. Regulatory mechanisms and sequence lengths in metalloregulator families.
- Figure S2. Distribution of pairwise sequence identity values across the 12 transcriptional regulator families
- Figure S3.
- Table S1. Summary of the information contained in the SSNs by metalloregulator family

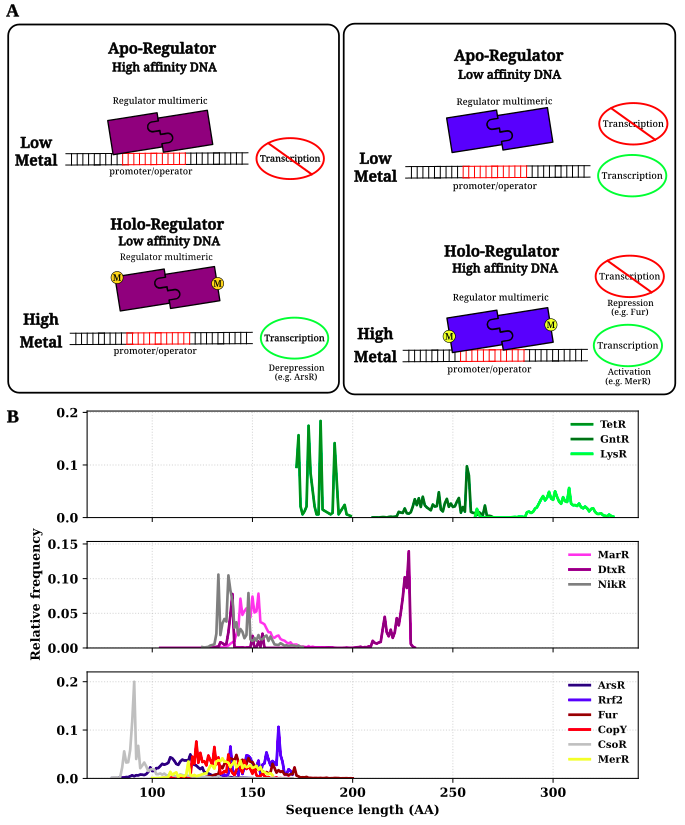

**Figure S1**. Regulatory mechanisms and sequence lengths in metalloregulator families. **A.** Known regulatory mechanisms in families whose affinity for DNA in its apo and holo (metalated) forms changes under different metal availability scenarios and can inhibit or activate downstream gene transcription. **B.** Sequence length distribution in different families of metalloregulators, separated into three main groups based on their folding and size. i) the compact and smaller ones ArsR, Rrf2, Fur, CopY, MerR, and CsoR; the medium-sized ones such as MarR, DtxR, and NikR; the largest ones such as TetR, GntR, and LysR.

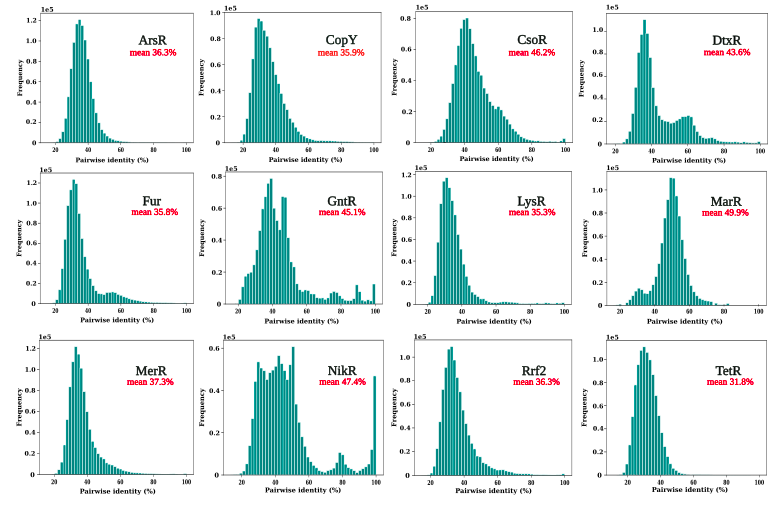

**Fig. S2.** Distribution of pairwise sequence identity values across the 12 transcriptional regulator families. For the TetR family, sequence identity values were calculated using a dataset filtered by sequence length (170–200 amino acids).

Table S1. Summary of the information contained in the SSNs by metalloregulator family

[**Click here**](https://docs.google.com/spreadsheets/d/1kKlRWJqY-x2CE-XPPMsuAsihBdd9zOakoA8g9H2lYHA/edit?usp=sharing)
